# An ODE-based mixed modelling approach for B- and T-cell dynamics induced by Varicella-Zoster Virus vaccines in adults shows higher T-cell proliferation with Shingrix compared to Varilrix

**DOI:** 10.1101/348284

**Authors:** Nina Keersmaekers, Benson Ogunjimi, Pierre Van Damme, Philippe Beutels, Niel Hens

## Abstract

Clinical trials covering the immunogenicity of a vaccine aim to study the longitudinal dynamics of certain immune cells after vaccination. The corresponding immunogenicity datasets are mainly analyzed by the use of statistical (mixed effects) models. This paper proposes the use of mathematical ordinary differential equation (ODE) models, combined with a mixed effects approach. ODE models are capable of translating underlying immunological post vaccination processes into mathematical formulas thereby enabling a testable data analysis. Mixed models include both population-averaged parameters (fixed effects) and individual-specific parameters (random effects) for dealing with inter-and intra-individual variability, respectively.

This paper models B-cell and T-cell datasets of a phase I/II, open-label, randomized, parallel-group study in which the immunogenicity of a new Herpes Zoster vaccine (Shingrix) is compared with the original Varicella Zoster Virus vaccine (Varilrix).

Since few significant correlations were assessed between the B-cell datasets and T-cell datasets, each dataset was modeled separately. By following a general approach to both the formulation of several different models and the procedure of selecting the most suitable model, we were able propose a mathematical ODE mixed-effects model for each dataset. As such, the use of ODE-based mixed effects models offers a suitable framework for handling longitudinal vaccine immunogenicity data. Moreover, it is possible to test differences in immunological processes between the two vaccines.

We found that the Shingrix vaccination schedule led to a more pronounced proliferation of T-cells, without a difference in T-cell decay rate compared to the Varilrix vaccination schedule.

**Author summary:** Upon vaccination, B-cells and T-cells are activated to induce an immune response against the vaccine antigen at hand. In this paper, we study and compare the longitudinal dynamics of the specific immune response based on a vaccine trial in which the immunogenicity of a new Herpes Zoster vaccine (Shingrix) is compared with the original Varicella Zoster Virus vaccine (Varilrix). We combine the use of ordinary differential equations (ODEs), i.e. mathematical models which are used to describe the dynamics of the immune response, with advanced regression analyses enabling us to infer the model parameters describing these dynamics. The resulting ODE-based mixed effects models enable describing the immune response dynamics allowing for both inter-and intra-individual variability; comparing the dynamics induced by the two vaccines and studying the B-and T-cell interactions. We found a more pronounced proliferation of T-cells for the Shingrix vaccination schedule as compared to the Varilrix vaccination schedule. The proposed methodology offers a suitable framework for better understanding the immunogenicity of vaccines.

## Introduction

Vaccines are developed in order to activate (and subsequently cause proliferation) of B-cells and T-cells that are specifically directed against the vaccine antigens. B-cells will (1) produce antigen-specific antibodies and (2) differentiate into long-living antibody secreting plasma cells. Antibodies are the primary effectors of the so-called humoral immune response in combating circulating pathogens. T-cells represent the cellular immune response and consist of CD4+ T-cells and CD8+ T-cells (and some other classes not discussed in this paper). CD4+ T-cells have an important role in helping other cell types (such as B-cells and macrophages) combating pathogens. CD8+ T-cells have a direct cytotoxic function and can target host cells that are infected by a pathogen.

The vaccine-induced B-cells and T-cells are hypothesized to be capable of preventing or minimizing the morbidity related to the infectious disease against which the vaccine is targeted. Vaccine immunogenicity trials aim to study the longitudinal dynamics of the specific immune response following vaccination. These trials can range from several months to several decades. The quantitative analysis of longitudinal immune response data has evolved from between-group and time point comparisons to statistical regression analyses [1, 2, 3]. Current state-of-the art statistical analyses of longitudinal data consist of a mixed effects model approach in which a separation is made between population-averaged parameters (so called fixed effects) and individual-specific parameters (so called random effects). More recently, Andraud et al. [4] and Le et al. [5] published the first papers in which the mixed effects modeling approach was combined with the use of ordinary differential equations (ODE), thereby more closely resembling immune response dynamics post vaccination. Whereas [4] focused on the long term dynamics following vaccination, [5] focused on the short term dynamics following vaccination.

ODE-based mathematical models are capable of translating the underlying immunological/biological theory into a testable data analysis. Moreover, the combination with mixed effects modeling offers a methodology capable of dealing with inter-and intra-individual variability. As such, the use of ODE-based mixed effects models offers a suitable framework for handling longitudinal vaccine immunogenicity data.

In this paper, we set out to use ODE-based mixed effects models to study B-cell and T-cell dynamics following varicella-zoster virus (VZV) vaccinations in VZV-immune adults. In particular, this framework will allow us to disentangle the immunogenic differences between two different VZV-specific vaccines.

In the Materials and methods section, we first present the immunogenicity data from two VZV vaccine studies consisting of B-cells and CD4+ T-cells of participants at different time points. We then describe the differential equations, the ODE and the ODE-based mixed effects models used to describe the immune response dynamics within each individual as well as the associated model selection procedures. In the Results section we apply the above methods and select a suitable model for each dataset. Next, we compare the results of the two VZV vaccines, using a group-related effect on a chosen parameter. Correlations in and between the datasets are also explored. Finally, we review our findings in the Discussion section.

## Materials and methods

### Data

The phase I/II, open-label, randomized, parallel-group study BIO101501 studies the safety and immunogenicity of an adjuvanted recombinant glycoprotein E vaccine (“HZ/su”, GSK) for VZV, by comparing it with a live attenuated Oka strain VZV vaccine (“OKA”, Varilrix^©^, GSK). To evaluate safety prior to administration in older adults, two groups of young adults (18-30 years) were vaccinated with two vaccine doses two months apart. The first group (GROUP 1; sample size: *n*_1_ = 10) received one dose of HZ/su and one dose of OKA concomitantly at month 0 and month 2 (i.e. four doses in total), whereas the second group (GROUP 2; *n*_2_ = 10) received a dose of HZ/su both times (i.e. two doses in total).

After vaccine safety was confirmed, three groups of older adults (50-70 years) were vaccinated two months apart, one group (GROUP 3; *n*_3_ = 45) received twice a single dose of HZ/su, the second (GROUP 4; *n*_4_ = 45) twice a single dose of OKA and the last (GROUP 5; *n*_5_ = 45) twice two concomitant doses of HZ/su and OKA. So, all in all, 155 participants were divided over these 5 groups. The properties of each group are summarized in Table 1.

Safety and immunogenicity were assessed for all groups up to 12 months post-vaccination in the original study. In order to obtain long-term immunogenicity data on the newly proposed HZ/su vaccine, 23 individuals from the groups solely receiving HZ/su (i.e. GROUPS 2 and 3) were assessed up to 42 months post-vaccination in the extension studies: BIO109671 and BIO109674. We refer to [6] for a more in depth description of the design and results of these studies.

**Table 1.** Properties of the different groups in the VZV vaccine trial.

| GROUP | sample size | age | vaccine | schedule |
| --- | --- | --- | --- | --- |
| 1 | 10 | 18-30 | HZ/su + OKA | 1+1 dose mo. 0, 1+1 dose mo. 2 |
| 2 | 10 | 18-30 | HZ/su | 1 dose mo. 0, 1 dose mo. 2 |
| 3 | 45 | 50-70 | HZ/su | 1 dose mo. 0, 1 dose mo. 2 |
| 4 | 45 | 50-70 | OKA | 1 dose mo. 0, 1 dose mo. 2 |
| 5 | 45 | 50-70 | HZ/su + OKA | 1+1 dose mo. 0, 1+1 dose mo. 2 |

Shown are sample size, age, vaccine and vaccination schedule (mo.=month). Group 4 is defined as reference group.

### B-cell data

First, we used data on the number of IgG-secreting B-cells, provided by a B-cell ELISPOT assay, at baseline, and at 1 month and 12 months after receiving the first vaccine dose. Two tests were performed: the first used Varilrix^©^ (1/20x) as stimulus in the B-cell ELISPOT assay, the second used 100 *μ*l of gE (10 *μ*g/ml) as stimulus. This resulted in two datasets comprising the frequencies of either “total” VZV-specific memory B-cells or gE-specific memory B-cells per million of IgG-producing memory B-cells. The participants were split into 5 different groups, based on age and vaccine type (see Section 2.1), and their profiles relative to the IgG-producing memory B-cells are plotted in Fig. 1 for the Varilrix-specific B-cells.

**Fig 1.**
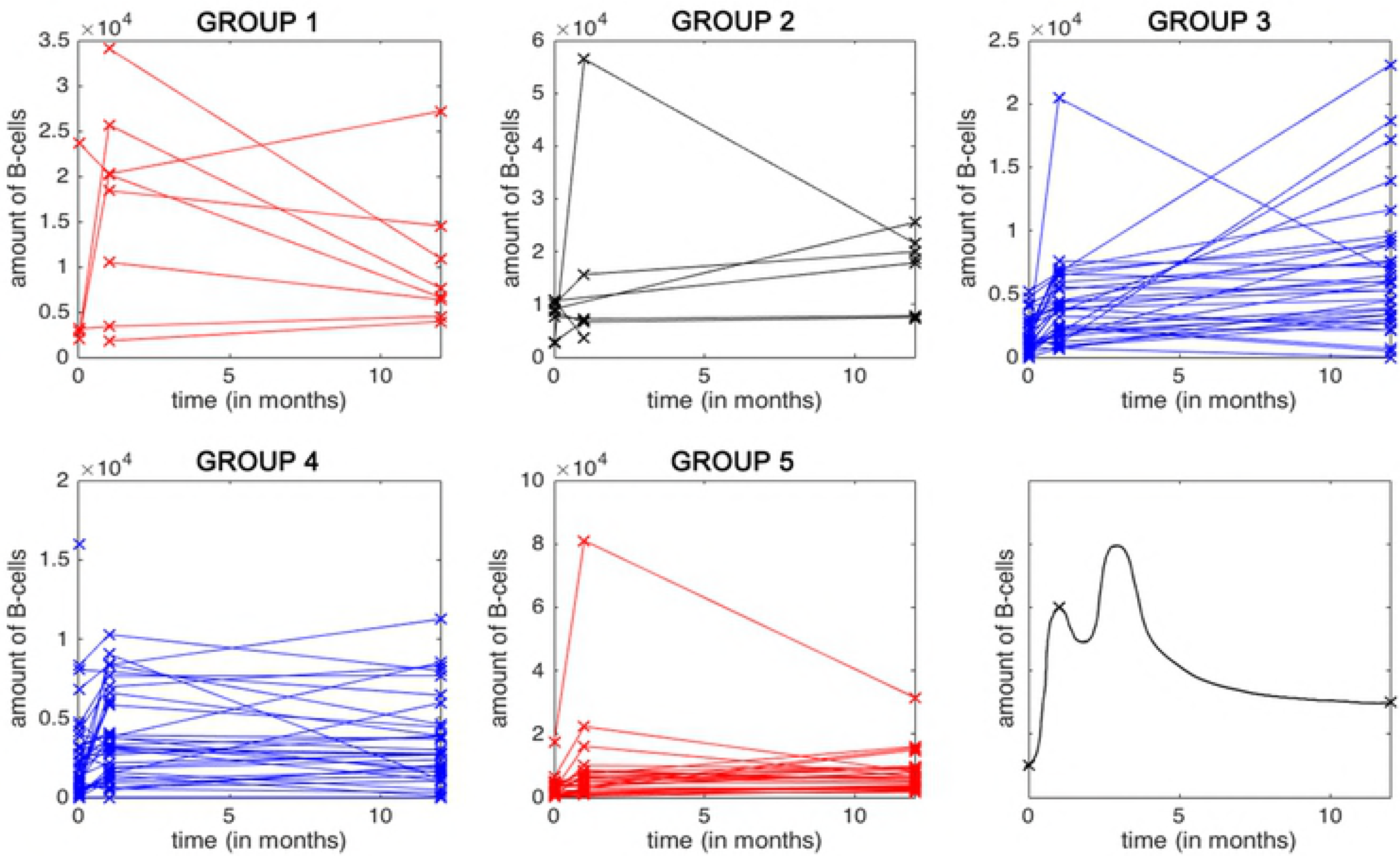
Amount of VZV IgG-secreting cells. Measured by B-cell ELISPOT (Varilrix stimulus) per 10^6^ IgG-secreting cells, up to 12 months. Data are shown per study group. The last panel shows a smooth function of the expected change in number of B-cells over time (in months), based on the observed data points per individual and considering the second vaccination at month 2.

We observe an increase in B-cells, or antibody-secreting cells (ASC), after vaccination at time *t* = 0 months. At time *t* = 2 months, the subjects were re-vaccinated, but no data were collected at that time point. Fig 1 shows only the time points for which data were available ( *t* = 0,1 and 12 months). Since it is reasonable to expect a (higher) peak in the data after the second dose at *t* = 2 months, we will assume a time period [0, *h*], *h* > 0 during which the level of B-cells increases up to a point, *h*, after which it decreases. The data plots of gE-specific ASC show a similar pattern (see Fig S1).

### T-cell data

Intracellular cytokine staining (ICS) in combination with a flow cytometric readout was performed to measure the amount of CD4+ T-cells that produced at least 2 cytokines (interferon-gamma, interleukin 2, CD40 Ligand, tumor necrosis factor alpha) using both Varilrix and gE as stimuli (in separate experiments). The subsequent two datasets comprise the same 155 participants, but now with time points at baseline, and at 1, 2, 3 and 12 months after receiving the first vaccine dose. The total VZV-specific T-cell profiles of the participants by study group are shown in Fig 2.

**Fig 2.**
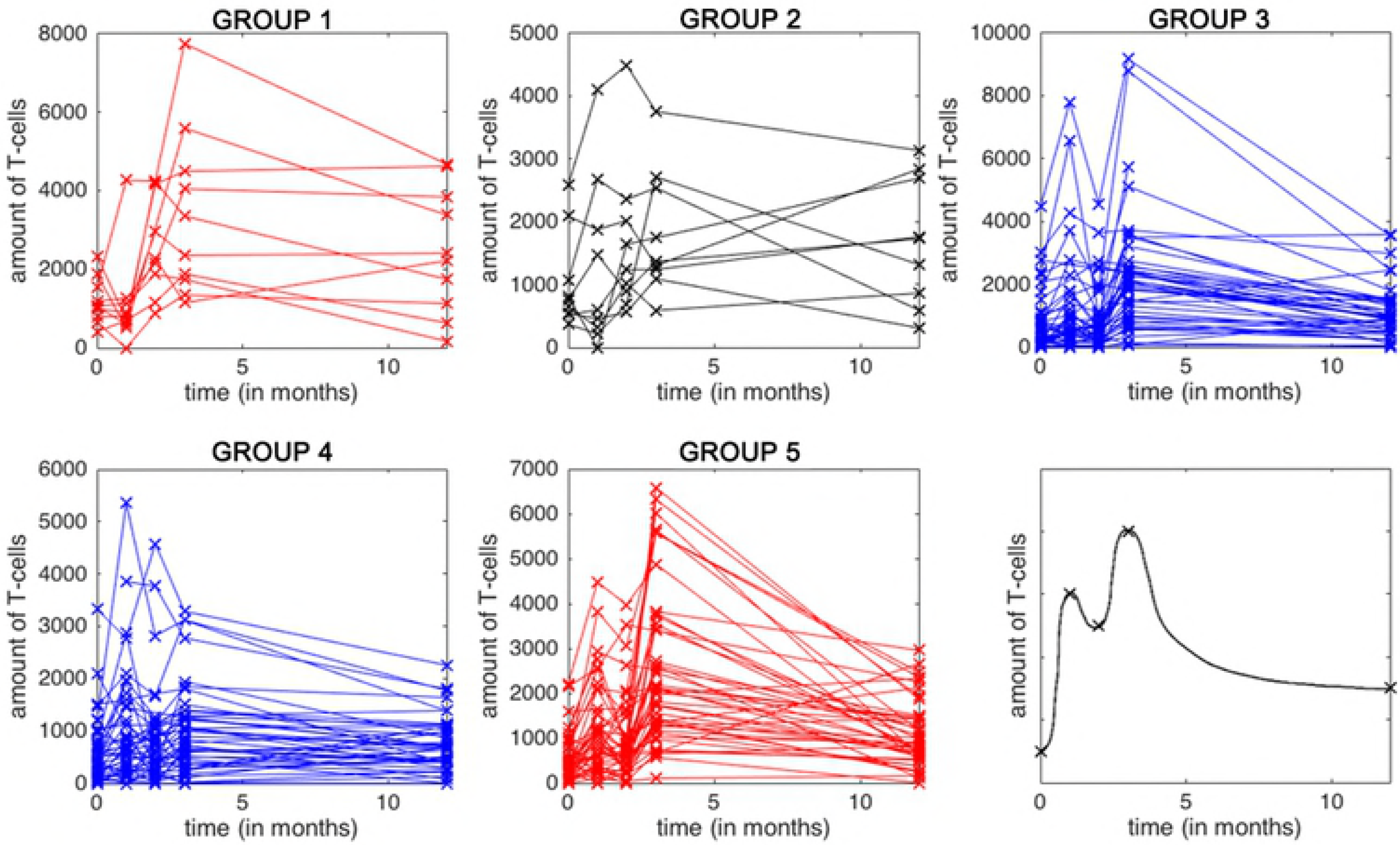
Amount of VZV-specific CD4+ T-cells, producing at least 2 immune markers. Measured by ICS per 10^6^ CD4+ T-cells, shown per group and up to 12 months. The last panel shows a smooth function of the expected change in number of T-cells over time (in months), based on the observed data points per individual.

Given that T-cell data were collected at more time points than B-cell data, we now observe a second peak in the group-specific data plots, as is expected given the vaccine administrations at month 2. Therefore we will use two time periods [0, *h*_1_] and [2, *h*_2_] (with 0 < *h*_1_ < 2 < *h*_2_) during which the level of T-cells first increases and then decreases. In case only one peak is observed, we will assume *h*_1_ = 2 < *h*_2_.

As with the B-cell profiles, the gE specific T-cell profiles are shown in Fig S2.

### Mathematical methods

We used systems of (nonlinear) ODEs to model the B-cell and T-cell dynamics. We applied a systematic approach to fit and compare several models in order to obtain the models that best describe the available data, while providing sufficient biological interpretation. The detailed version of all ODEs, along with their solution, can be found in S1 Appendix. In the following subsections we provide the basic rationale of these ODEs for both B-cell and T-cell dynamics, respectively.

### B-cell dynamics models

We describe the dynamics of the antibody secreting B-cells (ASC), using the following ODE:

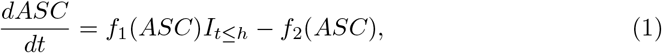

where *ASC*_0_ = *ASC*(0) denotes the initial number of antibody secreting cells at time=0 (months) and *f*_1_ and *f*_2_ are smooth functions of the number of ASC at time *t* (months), describing the change in the number of ASC due to activation and decay of ASC. We assume that the activation of ASC happens during a certain time period]0, *h*] and that after this time period, no new ASC are activated. The process of activation of ASC is described by the function *f*_1_. The decay in the number of ASC occurs at all times and is described by *f*_2_. In all models, the decay rate is assumed to remain constant over time.

A first distinction between models can be made through the nature of the rate at which ASC will be activated. A schematic overview of the different choices in activation/proliferation functions is given in Fig 3. In model B1, the proliferation rate is assumed to be constant over the time period [0, *h*[. Another option is to assume a rate which is proportional to the number of ASC at time *t*, as in model B2.

**Fig 3.**
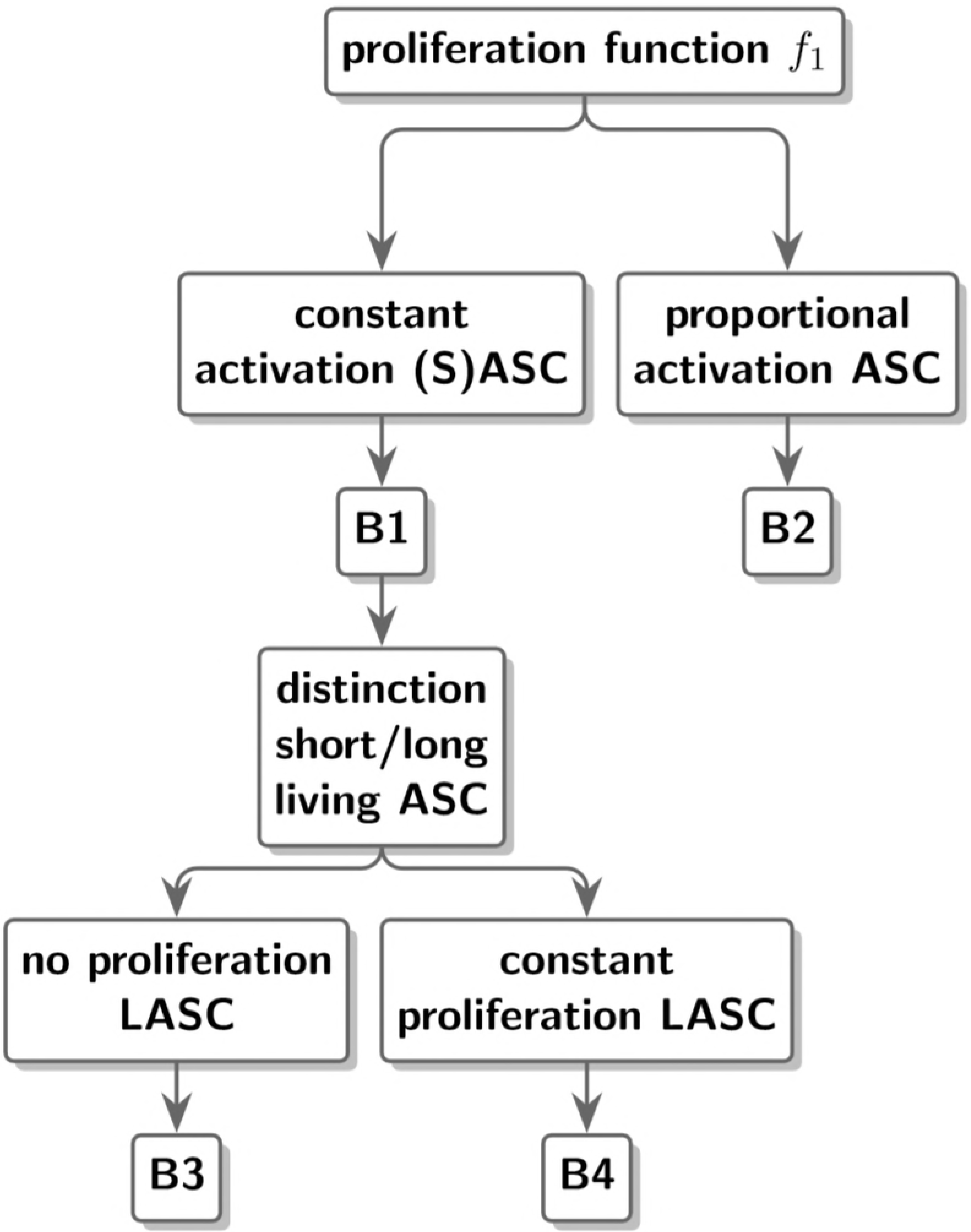
Schematic representation of the different proliferation functions used to describe the ASC population.

Following activation, B-cells can be divided into memory B-cells and plasma cells. Essentially, the former have a much shorter life span than the latter. Plasma cells remain in the bone marrow for a long time period [4, 7]. Models B3 and B4 incorporate this distinction by including different equations in the ODE system for short living ASC (SASC) and long living ASC (LASC). The dynamics of the SASC in models B3 and B4 are similar to those in model B1, but at time 0 months, no SASC are present in models B3 and B4. LASC however, are present in models B3 and B4 at time 0 months.

To distinguish between models B3 and B4, the dynamics of LASC are considered. First, in view of their long lifespan, no decay of LASC is assumed and model B3 expresses no proliferation rate either which means that the total number of LASC remains constant over time. Second, in model B4, a constant proliferation rate of LASC is introduced during time period [0, *h*[ after which their number will remain constant over time. Table 2 summarizes the parameters used in the functions of the different models.

**Table 2.** An overview of the different parameters in the B-cell dynamic models.

| Parameter | Description |
| --- | --- |
| $ASC_0$ | initial number of B-cells |
| $aASC$ | constant proliferation rate of ASC |
| $aSASC$ | constant proliferation rate of SASC (in case of a distinction) |
| $aLASC$ | constant proliferation rate of LASC (in case of a distinction) |
| $pASC$ | proportional proliferation rate of ASC |
| $uASC$ | decay of (short living, in case of a distinction) ASC |
| $h$ | time point after which no new ASC are activated |

### T-cell dynamic models

The design of the T-cell models follows a similar procedure as that of the B-cell models. The following ODE describes the basic dynamics of the stimulus-specific T-cell population:

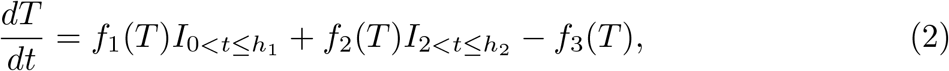

with *T*_0_ = *T*(0) the number of T-cells at time 0 (months). In this equation, *f*_1_(*T*) describes the proliferation of T-cells after the first vaccination event at time 0, which will occur until a certain time point *h*_1_ (with 0 < *h*_1_ ≤ 2). Afterwards, no T-cells will be activated until the second vaccination event 2 months after the first, which *f*_2_(*T*) describes as the proliferation of T-cells during the time period,]2, *h*_2_], with *h*_2_ the time point at which the second peak in T-cells is reached. The decay of T-cells will happen during the whole time period, and is represented by the function *f*_3_(*T*). In all models, it will be assumed that the decay rate of T-cells remains constant over time (cf. B-cell models). Moreover, we assume that the activation of T-cells after each vaccination event happens according to a constant proliferation rate. It is noteworthy that a non-constant proliferation rate, proportional to the number of T-cells, was part of exploratory analyses, but these explorations did not result in a convergent model (see the Inference and model selection section).

The rates after the first and second vaccination event are not necessarily equal and neither are the ranges of the time periods [0, *h*_1_[ and [2, *h*_2_[. Since T-cells are still present in the blood at the time of the second vaccination, we assume a different number of new T-cells will be activated.

Fig 4 summarizes the difference between all T-cell models we will consider. We start by assuming that all T-cells can be regarded as one population. With the additional assumptions that *f*_1_ and *f*_2_ are identical functions (and that the proliferation rates of T-cells are equal after each vaccination), we arrive at model T1. Model T2 does not presume both proliferation rates are equal.

**Fig 4.**
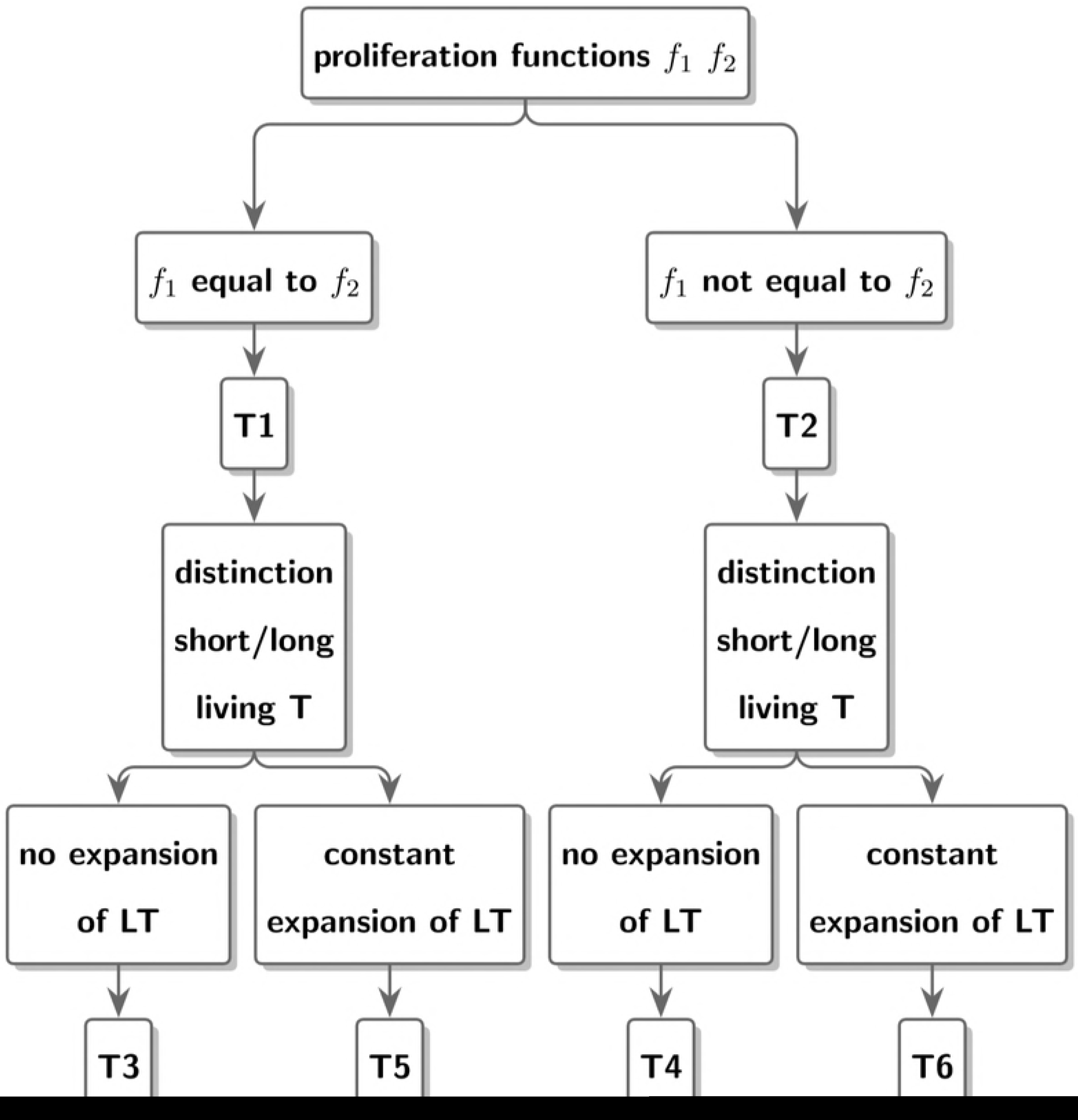
Schematic representation of the difference in proliferation functions used to describe the T-cell dynamics.

Next, models are considered in which the T-cell population is divided into short living and long living T-cell populations, represented by ST and LT, respectively. This can be considered as a distinction between effector T-cells (short living) and memory T-cells (long living).

The dynamics of the short living T-cells are similarly described as in models T1 and T2: a constant number of short living T-cells will be activated after each vaccination (not necessarily the same number), while the decay of short living T-cells occurs at all times at a constant decay rate. For models T3 and T4, the assumption is made that the total number of long living T-cells remains constant over time. If we add the distinction between models with equal and different functions *f*_1_ and *f*_2_, we arrive at models T3 and T4, respectively. Adding proliferation rates of long living T-cells after each vaccination, models T3 and T4 are extended to yield models T5 and T6. In order to restrict the total number of parameters, the long living T cell proliferation rates are assumed to be equal. Table 3 summarizes the parameters used in the functions of the different models.

**Table 3.** An overview of the different parameters in the T-cell dynamic models.

| Parameter | Description |
| --- | --- |
| $T_0$ | initial number of T-cells |
| $a_T$ | constant proliferation rate of T-cells after the $i$th vaccination |
| $a_{ST}$ | constant proliferation rate of short living T-cells (in case of a distinction) after $i$th vaccination |
| $a_{LT}$ | constant proliferation rate of long living T-cells (in case of a distinction) |
| $u_T$ | decay of (short living, in case of a distinction) T-cells |
| $h_i$ | time point after which no new T-cells are activated after the $i$th vaccination |

### Nonlinear mixed models

The dynamic models described in previous subsections can be formulated as nonlinear mixed models in which the parameters are assigned distributions through the specification of fixed and random effects. The fixed component can be interpreted as a population parameter, i.e. an average for all individuals, while the random component accounts for individual differences. More specifically, each individual parameter *P_i_* can thus be written as *P_i_* = *u_i_* × *P_pop_* where *P_pop_* is a population parameter and *u_i_* is log-normally distributed with E(*u_i_*) = 1.

In case of the presence of a categorical variable (e.g. different group in vaccine trial study), the different groups can be compared against each other by adding a component *β_j_* to the distribution of a certain parameter. *β_j_* describes how for group *j*, this parameter deviates from the (chosen) reference group. This makes it possible to test whether one group has a significant higher variable (e.g. rate of cell activation) compared to the reference group.

The Monolix software ^©^Lixoft was used for the estimation of the parameters. A built in stochastic approximation of the standard expectation maximization algorithm (SAEM) with simulated annealing, combined with a Markov Chain Monte Carlo (MCMC) procedure which replaces the simulation step of the SAEM algorithm, is used to obtain population parameters estimates. Loglikelihood calculation was done by importance sampling, in which a fixed *t*-distribution is assumed with 5 degrees of freedom. For more details on the algorithms used we refer to [8]. Mostly, Monolix default parameter values were used in the algorithms (see S2 Appendix.) The two-step SAEM-MCMC algorithm uses 10^6^ + 10^5^ iterations in order to assess convergence for estimating the population parameters.

### Inference and model selection

Although mathematical identifiability is guaranteed for the models presented in the Mathematical methods section, the complexity of these models when combining them with many random effects in view of the data limitations in terms of sampling times and sample sizes resulted in non-convergence. Therefore, simplifying assumptions needed to be made. One such simplifying assumption is presuming that the decay of B or T-cells is identical for all individuals, implying that the random effect for that decay parameter is omitted from the model.

For both the B-cell and T-cell data sets, the following procedure was used for comparing and selecting the most suitable biologically plausible model to describe the data. In a first step, a list of models was composed, consisting of models B1 to B4 for the B-cell data and of models T1 to T6 for T-cell data, together with assumptions on the parameters reflecting whether or not individual variation on these parameters is present, i.e. whether or not random effects were included for the different parameters. The model parameters were then estimated with the Monolix software.

Models with poor SAEM convergence, likely because of abundant model complexity, were discarded. Next, the candidate models were compared using Akaike’s Information Criterion (AIC) and the model with lowest AIC value was selected as first candidate model. Subsequently, a non-parametric bootstrap, using 1000 bootstrap re-samples, was performed on the candidate model. Since a sequential approach based on the candidate models with the lowest AIC values was used, the need to perform bootstraps for all candidate models was avoided, in order to decrease the number of computations. It was found that for a bootstrap, either 65%-77% of the samples had proper SAEM convergence, or the proportion of bootstrap samples with proper convergence was less than 15%. For this reason the criterion for good bootstrap convergence was defined as having at least 65% of bootstrap samples with proper SAEM convergence. In case of poor bootstrap convergence, the candidate model was rejected from the list of candidate models.

Next, a sensitivity analysis on the bootstrap results of the (converging) candidate model was performed by investigating whether the presence or absence of certain profiles of individuals in the bootstrap samples, had influence on its model convergence. For instance, if a single participant’s profile was more frequently part of non-converging datasets, a new bootstrap was performed, excluding the specific participant’s profile. Again, in case of poor bootstrap convergence the candidate model was rejected. If convergence remained sufficiently robust, it was investigated whether it was possible to improve the biological plausibility of the candidate model. This was done by examining the assumptions made on the parameters and adjusting those. An example of this is shown in the Model selection of T-cell datasets results section. The new model then became a candidate model.

### B-cell and T-cell dynamics associations

As CD4+ T-cells may directly influence the activation of B-cells, we investigated the existence of associations between B-cell and T-cell dynamics. We used a raw data complete cases analysis given that constructing a joint model based on the same concepts was not successful, likely due to data limitations (Inference and model selection section).

Since the process of B-cell activation is dependent on certain cytokine-expressing T-cells (such as CD40L), the hypothesis is investigated whether the increase in B-cells is proportional to the increase in T-cells. In order to examine this, we define *T*01:= *T*(1) - *T*(0) and *B*01:= *B*(1) - *B*(0) and use the `minerva` package in R to calculate the maximal information coefficient (MIC). This is a way to detect linear and non-linear relations between variables, and can thereby be used to indicate whether a linear relation is feasible by comparing it with the R-squared value [9]. In addition we compute the Spearman correlation.

Instead of solely examining the increase in B-cells and T-cells, we also looked at the correlation between the specific values of B-and T-cells at time points 0 and 1; Spearman correlations for all individuals were calculated between the data points B(0), B(1), T(0) and T(1).

## Results

### Model selection of B-cell datasets

We started by modeling the Varilrix-specific B-cell dataset, for which the model selection procedure outlined in the Inference and model selection section was followed. The upper part of Table 4 summarizes the differences between all models that were tested.

**Table 4.**
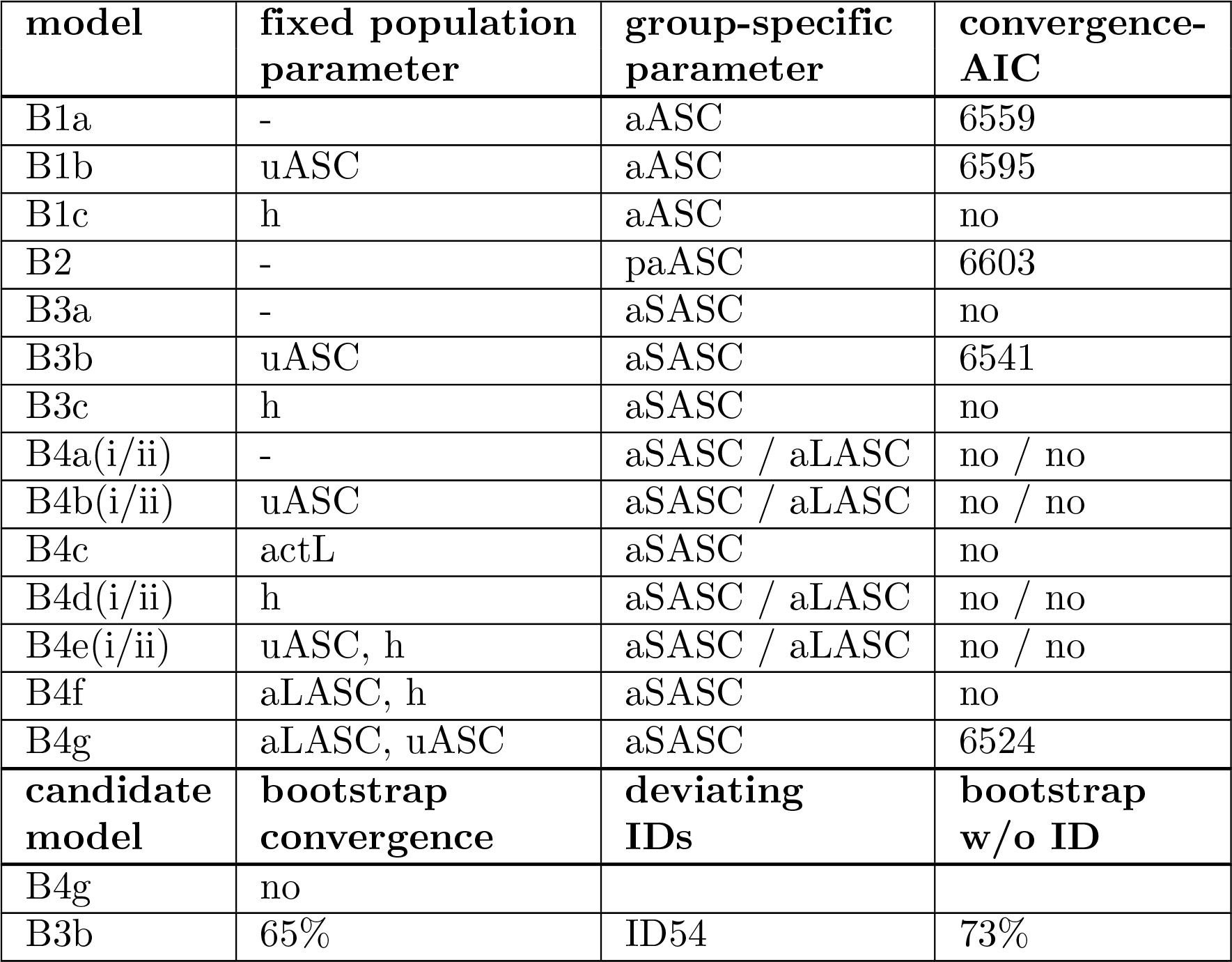
ODE Model formulations considered for Varilrix-specific B-cell data and model selection procedure.

| model | fixed population parameter | group-specific parameter | convergence-AIC |
| --- | --- | --- | --- |
| B1a | - | aASC | 6559 |
| B1b | uASC | aASC | 6595 |
| B1c | h | aASC | no |
| B2 | - | paASC | 6603 |
| B3a | - | aSASC | no |
| B3b | uASC | aSASC | 6541 |
| B3c | h | aSASC | no |
| B4a(i/ii) | - | aSASC / aLASC | no / no |
| B4b(i/ii) | uASC | aSASC / aLASC | no / no |
| B4c | actL | aSASC | no |
| B4d(i/ii) | h | aSASC / aLASC | no / no |
| B4e(i/ii) | uASC, h | aSASC / aLASC | no / no |
| B4f | aLASC, h | aSASC | no |
| B4g | aLASC, uASC | aSASC | 6524 |
| candidate model | bootstrap convergence | deviating IDs | bootstrap w/o ID |
| B4g | no |  |  |
| B3b | 65% | ID54 | 73% |

Upper part: Overview of the different models used to model the Varilrix-specific B-cell data. First column: model identifier. Second column: parameters selected as fixed population parameter. Third column: parameter which is chosen to be group-specific. Fourth column: AIC value of each model, in case of convergence.

Lower part: Overview of the considered candidate models (first column) used in the Varilrix-specific B-cell data model selection procedure. Second column: convergence results of the performed bootstraps. Third column: the results of possible IDs with deviating presence in the converging bootstrap samples. Fourth column: results of a bootstrap performed on the Varilrix-specific B-cell dataset, in case IDs are found.

For model B1, we distinguished between the following scenarios: a scenario with random effects for all parameters (model B1a), a scenario in which the decay of ASC (uASC) was assumed having a fixed population parameter (model B1b), and a scenario in which the time period where ASCs were activated (h) was fixed (model B1c). SAEM convergence was obtained for models B1a and B1b, with an AIC value of 6559 and 6595, respectively. Model B1c did not converge within 10^6^ + 10^5^ iterations and was therefore not considered any further.

Next, a proportional proliferation rate was explored in model B2. Model B2 assumed a group specific proliferation parameter *paASC* and all parameters having random effects. For this model, an AIC value of 6603 was obtained.

Consequently, we looked at models in which a distinction between SASC and LASC was made. Model B3a assumed all parameters had random effects. In models B3b and B3c, uASC and h, respectively, were set as fixed population parameter. Model B3b was the only model that showed convergence, with an AIC value of 6541.

The last model examined was model B4, in which a proliferation rate for LASC was added. Many assumptions on the parameters were made; the decay rate of ASC (uASC), proliferation rate of LASC (aLASC) and activation period (h) were set as fixed parameters in models B4b, B4c and B4d, respectively. Combinations of these fixed parameters were considered as well in models B4f, B4g and B4h. Apart from this, we also looked at different group specific parameters, not only the proliferation rate of SASC (aSASC) was considered, but the proliferation rate of long living B-cells (aLASC as well. Model B4g was the only model that accounted for aSASC and aLASC and still converged. In this model, both aLASC and uASC were set as fixed population parameters, and aSASC was considered to be a group specific parameter. This model had the lowest AIC value of 6524 among all aforementioned models, and was selected as first candidate model.

The bootstrap that subsequently was performed did not converge for model B4g, and it was therefore in the end rejected. Model B3b was selected as next-candidate model. The converging of its bootstrap was successful, with 65% of bootstrap samples showing proper SAEM convergence. An analysis was done to explore whether the presence (or absence) of certain individuals was responsible for the convergence of the datasets. When examining percentages of presence, ID54 showed deviant behavior: the profile was absent from a significant number of the non-converging datasets. It was therefore removed from the B-cell dataset, after which a new bootstrap with the candidate model was performed. Bootstrap convergence remained well (73 % of bootstrap samples showed SAEM convergence) and model B3b was therefore selected as final Varilrix-specific B-cell model. Model B3b differentiates between SASC and LASC LASC are assumed to remain constant through time. In time period (0, *h*), a constant number of SASC is activated and this proliferation rate is considered a group-specific parameter. The decay rate is assumed to be equal for each individual.

This model selection procedure is summarized in the lower part of Table 4. The estimated population parameters of the selected B-cell model are shown in Table 5.

**Table 5.** Varilrix-specific B-cell results.

| parameter | estimate | 95% CI |
| --- | --- | --- |
| $LASC_0$ | 1852.84 | (1425.79, 2407.80) |
| $aSASC$ | 774.17 | (367.67, 1630.12) |
| $\beta_1$ | 2.14 | (0.79, 3.49) |
| $\beta_2$ | 1.44 | (-0.02, 2.91) |
| $\beta_3$ | 0.67 | (-0.12, 1.46) |
| $\beta_5$ | 0.90 | (0.02, 1.79) |
| $uSASC$ | 0.13 | (0.11, 0.16) |
| $h$ | 3.30 | (2.05, 5.30) |

Parameter estimates and corresponding 95% confidence intervals (CI) of final model B3b. *SASC*(0) is assumed to be zero, *LASC*_0_ = *LASC*(0) denotes the initial number of LASC. The proliferation of SASC is constant in time period (0, *h*), at rate *aSASC* and assumed to be group-specific with effects *β*_*i*_ (*i* = 1, 2,3, 5). Decay of SASC happens at rate *uSASC*. The number of LASC remains constant through time.

The same models as with the Varilrix-specific B-cell data were used for the gE-specific B-cell data and a similar model selection procedure was followed. For more details, we refer to S3 Appendix. The outcome of the model selection, was model B1a, a model which does not differentiate between SASC and LASC. In time period (0, *h*), a constant number of ASC are activated. All parameters are assumed to have random effects and the activation rate of ASC is chosen as group-specific parameter. The parameter estimations, along with confidence interval, are shown in Table 6.

**Table 6.**
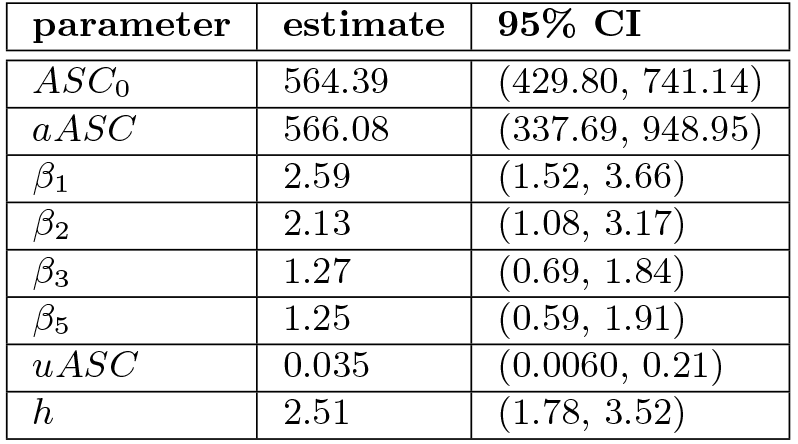
gE-specific B-cell results.

| parameter | estimate | 95% CI |
| --- | --- | --- |
| $ASC_0$ | 564.39 | (429.80, 741.14) |
| $aASC$ | 566.08 | (337.69, 948.95) |
| $\beta_1$ | 2.59 | (1.52, 3.66) |
| $\beta_2$ | 2.13 | (1.08, 3.17) |
| $\beta_3$ | 1.27 | (0.69, 1.84) |
| $\beta_5$ | 1.25 | (0.59, 1.91) |
| $uASC$ | 0.035 | (0.0060, 0.21) |
| $h$ | 2.51 | (1.78, 3.52) |

Parameter estimates and corresponding 95% confidence intervals (CI) of final model B1a. No distinction between SASC and LASC is presumed. *ASC*_0_ = *ASC*(0) denotes the initial number of ASC. The proliferation of ASC is constant in time period (0, *h*), at rate *aASC* and assumed to be group-specific with effects *β_i_* (*i* = 1, 2, 3, 5). Decay of ASC happens at rate *uSASC*.

### Model selection of T-cell datasets

Just like with the B-cell models, the most simplistic T-cell model T1 was considered first, in which *a*_1_*T* = *a*_2_*T* = *aT*. Model T1a assumed all parameters had random effects, and the activation of T-cells (*aT*) had a group-specific effect. An AIC value of 11,664 was obtained.

When assuming *a*_1_*T* ≠ *a_2_T*, we arrived at model T2. First, the assumption was made that all parameters had random effects, and both *a*_1_*T* and *a*_2_*T* had a group-specific effect in model T2a. This model converged, with an AIC value of 11,658. When assuming only *a*_2_*T* had a group-specific effect (model T2b), a slightly lower AIC value was obtained at 11,655.

Next, a distinction between short and long living T-cells (ST and LT, respectively) was considered. In model T3a, all parameters had random effects and *aST* (= *a*_1_*ST* = *a*_2_*ST*) had a group-related effect resulting in an AIC value of 11,637. Model T3b, in which *uT* was a fixed parameter, did not reach convergence.

Model T4 assumed different activation rates of T-cells after each vaccination. When all parameters had random effects, and both *a*_1_*T* and *a*_2_*T* were group specific, SAEM convergence was not reached (model T4a). When only a group specific effect on *a*_2_*T* was assumed (model T4b), convergence was achieved resulting in an AIC value of 11,615. Subsequently, *uT* was set as fixed parameter in models T4c and T4d, again with group specific effects on both activation rates (T4c) and on *a*_2_*T* only (T4d), respectively. Both models showed SAEM convergence with an AIC value of 11,646 and a lower AIC value of 11,626, respectively.

When assuming LT activation according to a constant proliferation rate (equal after each vaccination in order to limit the number of parameters to be estimated), models T5 and T6 were reached. In models T5, the activation rates of ST were presumed equal after each vaccination. Together with the assumption that all parameters were random, and *aST* was a group specific parameter, this leaded to model T5a, where an AIC value of 11,631 was found. We note that setting *aLT* as a group specific parameter was tested as well, but none of these models (including the following) showed convergence and thus were omitted from Table 7.

**Table 7.**
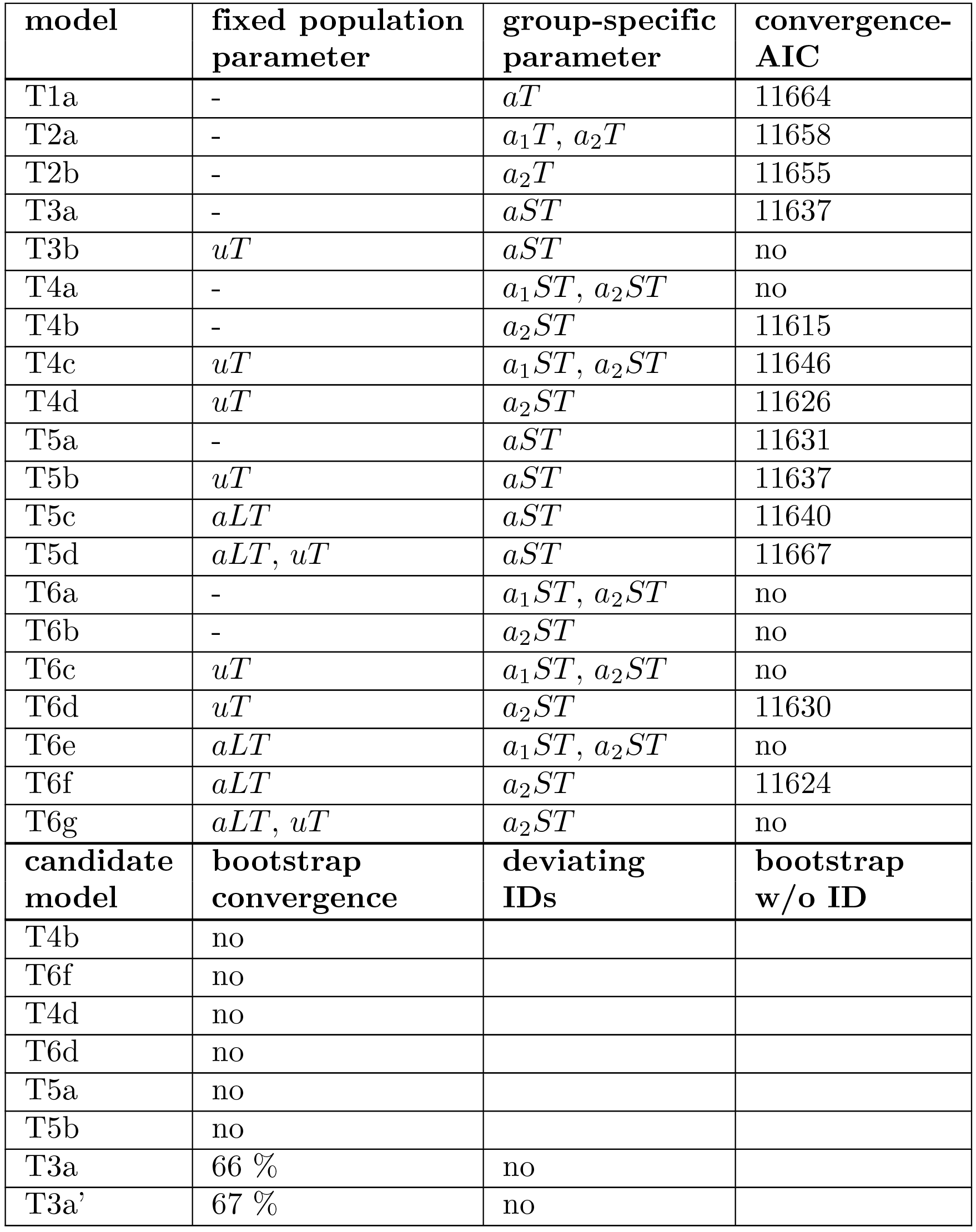
ODE Model formulations considered for Varilrix-specific T-cell data and model selection procedure.

| model | fixed population parameter | group-specific parameter | convergence-AIC |
| --- | --- | --- | --- |
| T1a | - | $aT$ | 11664 |
| T2a | - | $a_1T, a_2T$ | 11658 |
| T2b | - | $a_2T$ | 11655 |
| T3a | - | $aST$ | 11637 |
| T3b | $uT$ | $aST$ | no |
| T4a | - | $a_1ST, a_2ST$ | no |
| T4b | - | $a_2ST$ | 11615 |
| T4c | $uT$ | $a_1ST, a_2ST$ | 11646 |
| T4d | $uT$ | $a_2ST$ | 11626 |
| T5a | - | $aST$ | 11631 |
| T5b | $uT$ | $aST$ | 11637 |
| T5c | $aLT$ | $aST$ | 11640 |
| T5d | $aLT, uT$ | $aST$ | 11667 |
| T6a | - | $a_1ST, a_2ST$ | no |
| T6b | - | $a_2ST$ | no |
| T6c | $uT$ | $a_1ST, a_2ST$ | no |
| T6d | $uT$ | $a_2ST$ | 11630 |
| T6e | $aLT$ | $a_1ST, a_2ST$ | no |
| T6f | $aLT$ | $a_2ST$ | 11624 |
| T6g | $aLT, uT$ | $a_2ST$ | no |
| candidate model | bootstrap convergence | deviating IDs | bootstrap w/o ID |
| T4b | no |  |  |
| T6f | no |  |  |
| T4d | no |  |  |
| T6d | no |  |  |
| T5a | no |  |  |
| T5b | no |  |  |
| T3a | 66 % | no |  |
| T3a' | 67 % | no |  |

Above: Overview of the different models used to model the Varilrix-specific T-cell data. First column: model identifier. Second column: parameters selected as fixed population parameter. Third column: parameter which is chosen to be group-specific. Fourth column: AIC value of each model, in case of convergence.

Under: Overview of the considered candidate models (first column) used in the Varilrix-specific T-cell data model selection procedure. Second column: convergence results of the performed bootstraps. Third column: the results of possible IDs with deviating presence in the converging bootstrap samples. Fourth column: results of a bootstrap performed on the Varilrix-specific T-cell dataset, in case IDs are found.

In Models T5b and T5c, respectively, *uT* and *aLT* were assumed to be fixed population parameters. They showed SAEM convergence, with AIC values of 11,637 and 11,640, respectively. Setting both *uT* and *aLT* fixed in model T5d did not improve the model (AIC: 11,667).

Finally, model T6 was considered, with different activation rates after each vaccination event. Assuming all parameters were random and either both *a*_1_*ST* and *a*_2_*ST* (T6a), or only *a*_2_*ST* (T6b), were group specific, did not lead to convergence within 10^6^ + 10^5^ iterations. For this reason, fixed parameters *uT* and/or *aLT* were considered. When *uT* was fixed, and both activation rates were group specific, convergence was not reached (model T6c). With *a*_2_*ST* being group specific (model T6d), convergence was obtained with an AIC value of 11,630. In case of setting *aLT* as a fixed parameter, similar results were found; assuming both activation rates to be group specific did not lead to convergence, but assuming only *a*_2_*ST* was group specific, did, with a slightly lower AIC value equal to 11,624. The last scenario assumed both *aLT* and *uT* were fixed population parameters, though no convergence was obtained.

Since model T4b was the model with lowest AIC (11,615), it was selected as first candidate model. However, model T4b was subsequently rejected as a 1000 sample bootstrap failed to converge. Likewise models T6f, T4d, T6d, T5a and T5b did not have proper bootstrap convergence. Next, model T3a was selected as candidate model and showed bootstrap convergence; 66% of the bootstrap samples reached SAEM convergence. As before, a search for frequently deviant profiles in the converging and non-converging bootstrap datasets was performed, but no such profile was identified.

Taking into account that the assumption that *a*_1_*ST* = *a*_2_*ST* = *aST* might not be a realistic assumption in a model that described a real life cellular process, a difference in proliferation rates after each vaccination was inserted, assuming *a*_2_*ST* was proportional to *a*_1_*ST*: *a*_2_*ST* = *k* × *a*_1_*ST*. Adding this parameter to the pool of parameters to be estimated in the SAEM algorithm, did not yield a converging model. In order to limit the number of parameters to be estimated by the SAEM-algorithm, different values of *k* ware set fixed and for each subsequent model, AIC values were compared. A model with *k* = 0.15 showed the lowest AIC value (11,623), which we named model T3a’, and a bootstrap with 100 bootstrap samples was performed on this model. From this bootstrap, 67% of samples resulted in SAEM convergence and a search for disproportionate presence of deviant profile(s) did not identify such profiles. Therefore, model T3a’, a model which differentiates between ST and LT, was selected as final Varilrix-specific T-cell model. This model assumes the number of LT remains constant through time and assumes activation of ST is constant in time periods (0, *h*_1_) and (2, *h*_2_) with *a*_2_*ST* = 0.15 × *a*_1_*ST*. Moreover, all parameters are assumed to have random effects with the activation rate a group-specific parameter. The model parameter estimates are shown in Table 8.

**Table 8.**
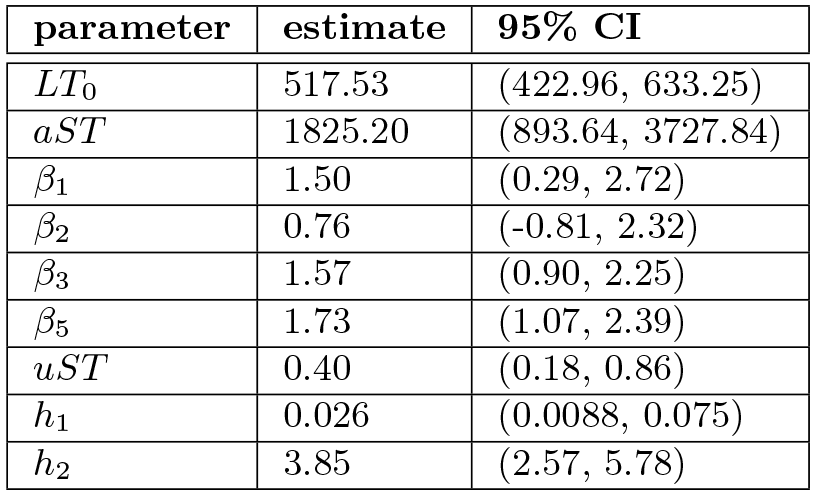
Varilrix-specific T-cell results.

Parameter estimates and corresponding 95% confidence intervals (CI) of final model T3a’. *ST*(0) is assumed to be zero, *LT*_0_ = *LT*(0) denotes the initial number of LT. The proliferation of ST is constant in time period (0, *h*_1_) at rate *aST* and in time period”2, *h*_2_) at rate 0.15.*aST*. *aST* is assumed to be group-specific with effects *β_i_* (*i* = 1, 2, 3, 5). Decay of ST happens at rate *uST*. The number of LT remains constant through time.

The same T-cell models were used in the model selection procedure of the gE-specific T-cell data, more details are found in S3 Appendix. The parameter estimations of the final gE-specific T-cell model T1a” are shown in Table 9. Model T1a” does not differentiate between ST and LT and assumes a constant activation of T-cells in time periods (0, *h*_1_) and (2, *h*_2_) with *a*_2_*ST* = 0.66 × *a*_1_*ST*. All parameters were assumed to have random effects with the activation rate being a group-specific parameter. Moreover, individuals 89 and 149 were left out of this dataset since they had too much influence on model convergence.

**Table 9.** gE-specific T-cell results.

| parameter | estimate | 95% CI |
| --- | --- | --- |
| $T_0$ | 90.07 | (45.31, 179.07) |
| $aT$ | 329.14 | (162.70, 665.81) |
| $\beta_1$ | 2.41 | (1.75, 3.32) |
| $\beta_2$ | 2.30 | (1.65, 3.21) |
| $\beta_3$ | 2.53 | (1.93, 3.31) |
| $\beta_5$ | 2.43 | (1.83, 3.22) |
| $uT$ | 0.54 | (0.26, 1.11) |
| $h_1$ | 0.21 | (0.15, 0.31) |
| $h_2$ | 9.03 | (5.61, 14.53) |

Parameter estimates and corresponding 95% confidence intervals (CI) of final model T1a’. No distinction of ST and LT is presumed. *T*_0_ = *T*(0) denotes the initial number of T-cells. The proliferation of T-cells is constant in time period (0, *h*_1_) at rate *aT* and in time period (2, *h*_2_) at rate 0.66.*aT*. *aT* is assumed to be group-specific with effects *β_i_* (*i* = 1, 2, 3, 5). Decay of T-cells happens at rate *uT*.

### Vaccine differences

Group-specific effects on chosen parameters make it possible to compare each group by examining the differences in these effects. This comparison focuses on different proliferations of B-and T-cells. Group-specific effects were also added on other parameters, more specifically the decay rate and time point *h*, which marks the end of the proliferation period after vaccination. However, models with these group-specific parameters did either not have an increased AIC compared to a model without this effect, or did not show SAEM convergence.

Since a group-specific component was added to the activation of B-/T-cells for each final model, it was subsequently possible to examine whether the HZ/su vaccine caused a higher increase in B-and or T-cells after vaccination, compared to the original OKA vaccine.

As a reminder, Table 1 summarizes the characteristics of the 5 different groups in the vaccine trial. Group 4 received the original OKA vaccine and was thereby defined as reference group.

Next, we calculated corresponding p-values of the group-specific parameters that were estimated in the Model selection of B-cell datasets and T-cell datasets sections, shown in Table 10. In view of the sample size of groups 1 and 2 (and age), we were mainly interested in *β*_3_ and β_5_. A *β_i_* higher than zero indicates a higher activation rate of cells in the groups receiving the HZ/su vaccine, compared to the activation rate in the reference group which received the OKA vaccine. In the case of Varilrix-and gE-specific T-cells, both groups 3 and 5 showed a significant higher activation rate (*p* < 0.05). The activation rate of gE-specific B-cells also was significantly higher compared to the reference group. Varilrix-specific B-cells also seemed to have a higher proliferation rate, though in the case of groups receiving solely the HZ/su vaccine, not significantly so (*p* > 0.05).

**Table 10.** Differences in proliferation rates per group.

|  | Varilrix B-cells | gE B-cells | Varilrix T-cells | gE T-cells |
| --- | --- | --- | --- | --- |
| $\beta_1$ (CI),<br>p-value | 2.14 (0.79, 3.49)<br>$2.8 \times 10^{-4}$ | 2.59 (1.52, 3.66)<br>$< 10^{-5}$ | 1.50 (0.29, 2.72)<br>$3.8 \times 10^{-4}$ | 2.41 (1.75, 3.32)<br>$< 10^{-5}$ |
| $\beta_2$ (CI),<br>p-value | 1.44 (-0.02, 2.91)<br>0.013 | 2.13 (1.08, 3.17)<br>$< 10^{-5}$ | 0.76 (-0.81, 2.32)<br>0.12 | 2.30 (1.65, 3.21)<br>$< 10^{-5}$ |
| $\beta_3$ (CI)<br>p-value | 0.67 (-0.12, 1.46)<br>0.10 | 1.27 (0.69, 1.84)<br>$10^{-5}$ | 1.57 (0.90, 2.25)<br>$< 10^{-5}$ | 2.53 (1.93, 3.31)<br>$< 10^{-5}$ |
| $\beta_5$ (CI)<br>p-value | 0.90 (0.02, 1.79)<br>0.025 | 1.25 (0.59, 1.91)<br>$3 \times 10^{-5}$ | 1.73 (1.07, 2.39)<br>$< 10^{-5}$ | 2.43 (1.83, 3.22)<br>$< 10^{-5}$ |

Group-specific parameter estimates, corresponding 95% confidence intervals (CI) and p-values, calculated for Varilrix B-cell, gE B-cell, Varilrix T-cell and gE T-cell data.

### Potential associations

We started by examining the Varilrix-specific B-cell and T-cell datasets. The datasets were restricted to individuals without missing values of B-cell or T-cell data at time points 0 and 1 month, 96 individuals in total. First, the hypothesis was made that an increase in T-cells was proportional to an increase in B-cells. We express the expected proportionality factor by *m*:

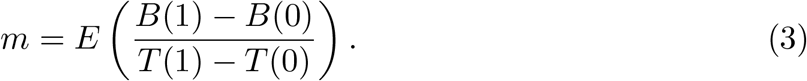

Fig 5 shows a scatterplot of *T*01:= *T*(1) − *T*(0) plotted against *B*01:= *B*(1) − *B*(0). At first sight, a linear relation between *T*01 and *B*01 might not seem a reasonable assumption.

**Fig 5.**
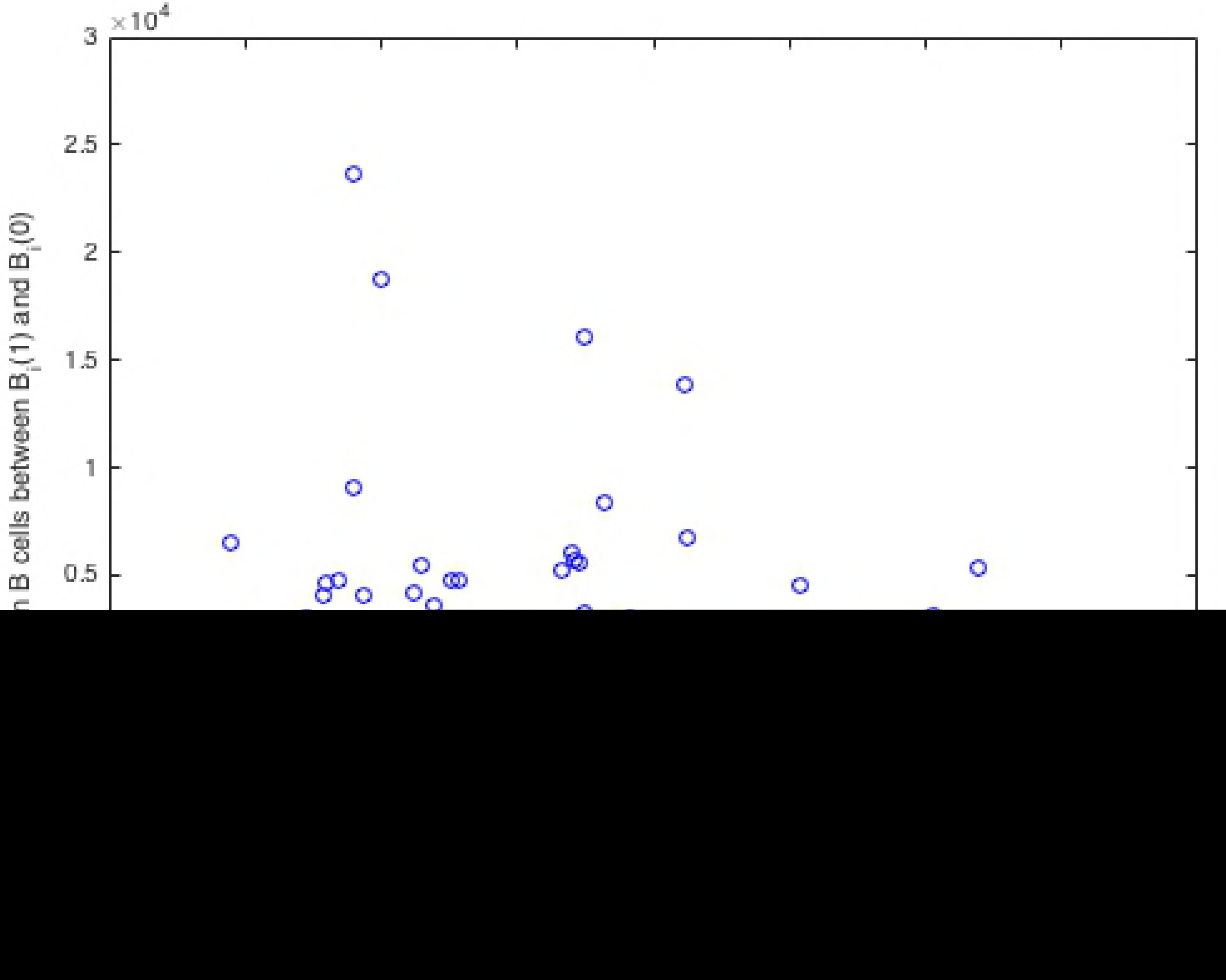
Scatterplot of the increase in Varilrix-specific T-cells (T01), plotted against the increase in Varilrix-specific B-cells (B01)

To further examine this, we calculated the Spearman correlation and the maximal information coefficient (MIC) as a way to assess and measure (non)linear relationships between datasets.

The Spearman correlation between Varilrix-specific *T*_01_ and *B*_01_ was −0.0274 (*p* = 0.7914), which rejected the hypothesis that increases in Varilrix-specific T-cells were associated to increases in Varilrix-specific B-cells. A non-significant MIC of 0.2161 (5% significance level) confirmed this result.

We also assessed MIC and Spearman correlations on the same datasets per subgroup, reaching the same conclusion (Table 11).

**Table 11.** Correlations between B-cells and T-cells (Varilrix stimulus)

| group | MIC | p-value | Spearman | p-value |
| --- | --- | --- | --- | --- |
| 3 | 0.2854 | $> 0.05$ | 0.2096 | 0.2663 |
| 4 | 0.1673 | $> 0.05$ | -0.2192 | 0.2271 |
| 5 | 0.2611 | $> 0.05$ | 0.1009 | 0.6380 |

MIC coefficients, Spearman correlation and corresponding p-values between increase in Varilrix-specific B-cells (*B*01) and increase in Varilrix-specific T-cells (*T*01), calculated for groups 3, 4 and 5. As the sample sizes of groups 1 and 2 were too small (*n*_1_ = 4 and *n*_2_ = 6), those groups were omitted from the analysis.

Potential associations between increases in gE-specific T-cells and B-cells were investigated. The scatterplot of *T*_01_ plotted against *B*_01_ is found in Fig S3.

The Spearman correlation of 0.3833 (*p* = 1.0647*e*^−04^) suggested there was indeed an association. The MIC score was calculated as 0.4107, which was significant (*p* < 0.001). In case of a linear relation, the R-square is expected to be close to this MIC score. As the R-square was equal to 0.05593, we could exclude a linear relationship.

As before, we also studied the relations between *T*_01_ and *B*_01_ per subgroup (see Table 12). The Spearman p-values showed that only in group 3 the Spearman correlation could be considered significant, together with MIC, implying a nonlinear relationship.

**Table 12.** Correlations between B-cells and T-cells (gE stimulus)

| group | MIC | p-value | MIC-$R^2$ | Spearman | p-value |
| --- | --- | --- | --- | --- | --- |
| 3 | 0.4734 | 0.01007 | 0.4665 | 0.4725 | 0.0084 |
| 4 | 0.3052 | > 0.05 |  | -0.2300 | 0.2053 |
| 5 | 0.3149 | > 0.05 |  | 0.3138 | 0.1266 |

MIC coefficients, Spearman correlation and corresponding p-values between increase in gE-specific B-cells (*B*01) and increase in gE-specific T-cells (*T*01), calculated for groups 3, 4 and 5. As the sample sizes of groups 1 and 2 were too small (*n*_1_ = 4 and *n*_2_ = 6), those groups were omitted from the analysis.

Correlations between the initial number of B-cells B(0), the initial number of T-cells T(0), the number of B-cells at month 1 B(1) and the number of T-cells at month 1 T(1) have also been investigated.

We started with the Varilrix-specific B-cell and T-cell data. Again, we only included individuals for whom we had data points B(0), B(1), T(0) and T(1). Spearman correlations for all individuals were calculated between these data points, and the results are shown in Fig 6. This figure also shows the Spearman correlations when we separated the individuals by group. When looking at the correlations of all individuals, we noticed correlations between B(0) and B(1), and T(0) and T(1), but no significant correlations between B-and T-cells. Examining the correlations by group, it seemed those differ greatly depending on the group. However, it has to be noted that group 1 and 2 contain a very small number of individuals (4 and 6 individuals respectively). For groups 3 to 5, the results were similar to the results of the correlation between all individuals, and we therefore found no convincing evidence for an association between Varilrix specific B-cells and T-cells.

**Fig 6.**
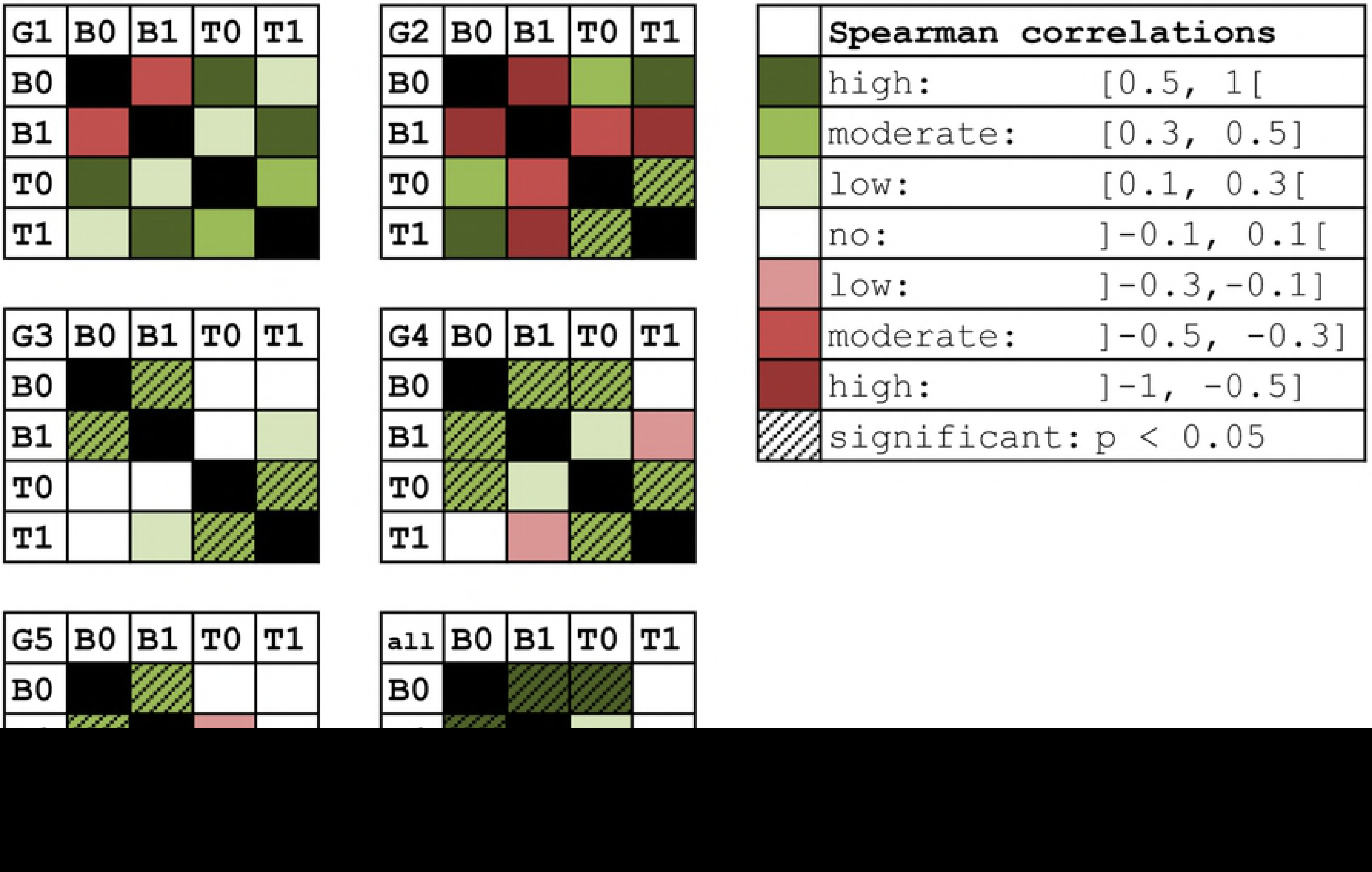
Spearman correlation between Varilrix-specific data points B(0), B(1), T(0) and T(1) Shown for all individuals and per group. Significant correlations are indicated.

The correlation between data points at time 0 and 1 of gE-specific B-cells and T-cells was examined next. Fig 7 shows the Spearman correlations, first for all individuals and then split by group. We observed that some correlations seemed to be higher compared to the Varilrix-specific data, however, when examining the Spearman matrices by group, the values between the different groups seemed to vary widely. For this reason, we did not find decisive evidence to include the number of gE-specific T-cells into the B-cell models or vice versa.

**Fig 7.**
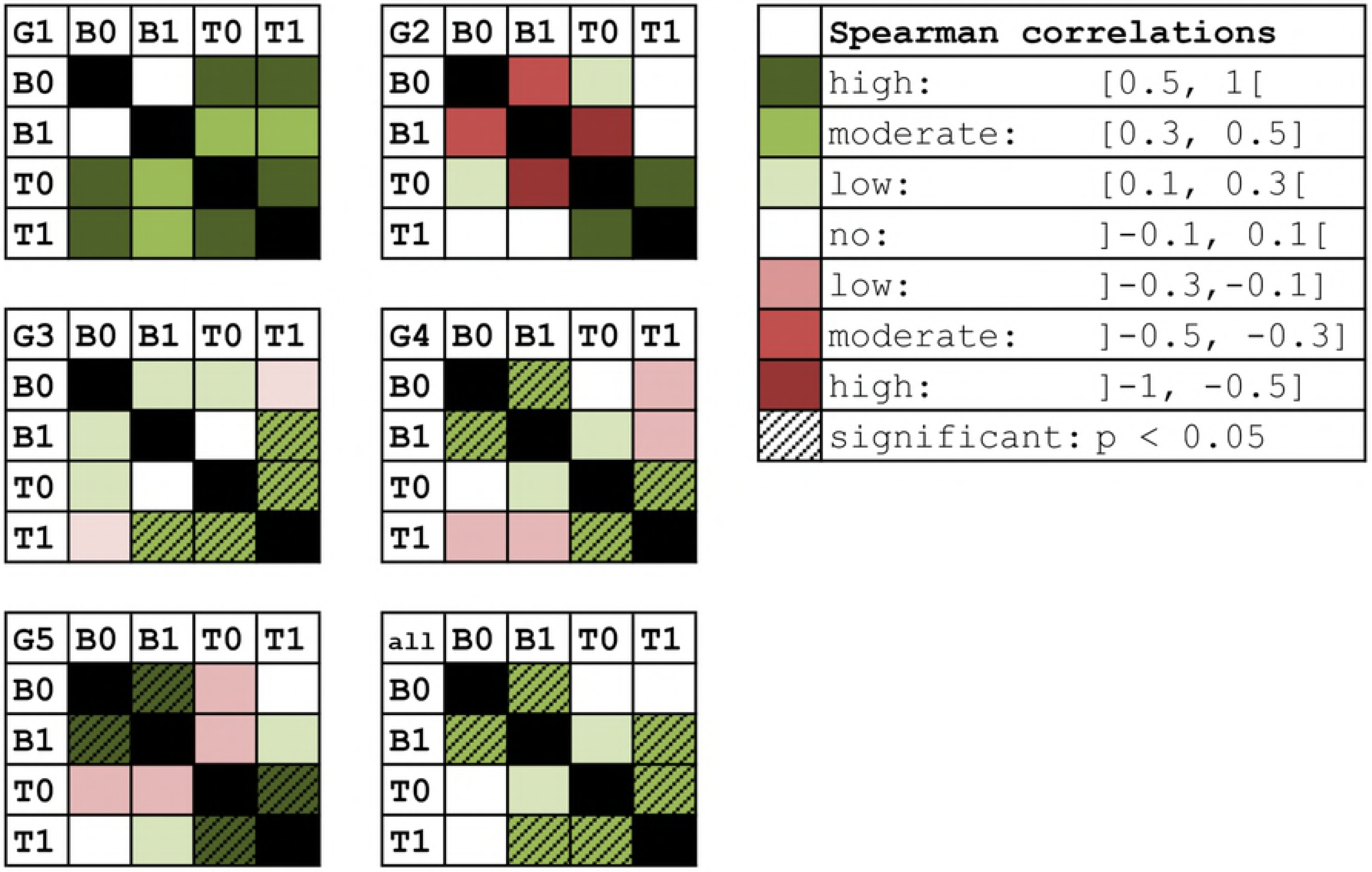
Spearman correlation between gE-specific data points B(0), B(1), T(0) and T(1) Shown for all individuals and per group. Significant correlations are indicated.

## Discussion

In this study we used a nonlinear mixed modeling approach using ordinary differential equations to describe B-cell and T-cell dynamics in adults following 2-dose vaccination against VZV by means of the novel subunit VZV gE vaccine (Shingrix, GSK) and live-attenuated VZV (Varilrix, GSK). Our study was motivated by the difficulties in attributing differences between vaccines and vaccination schedules to underlying immunological processes when using “classical” statistical techniques that do not take into account the underlying immunology. Recently, Andraud et al. [4] and Le et al. [5] showed that nonlinear ODE mixed modeling in the setting of vaccinations was capable of giving estimates on several biological parameters.

In our study, we assessed Varilrix and VZV gE-specific B-cell and T-cell responses during the 2-dose vaccination schedule for three different schedules (Shingrix only, Varilrix only and the combination of Shingrix and Varilrix on both vaccination moments). We used a standardised inference method to obtain for each setting (immune response and vaccine) the most optimal equation model and parameter estimation (using bootstrapping). Using this robust approach we were able to conclude that the best models did not have an overly complex structure. Models with constant proliferation rates (after each vaccination, in time periods (0, *h*) (B-cells) and (0, *h*_1_) and (2, *h*_2_) (T-cells)) had lower AIC compared to models with proportional proliferation rates. Restricting the number of parameters, either by not making a distinction between short and long living B-/T-cells (gE-specific B-and T-cell models B1a and T1a’), or by assuming the number of long living B-/T-cells remains constant through time (Varilrix-specific B-and T-cell models B3b and T3a’), proved to lead to preferable models compared to the other considered model structures. Although some of these other models may have had a more intuitively logical biological interpretation, they often did not have SAEM or bootstrap convergence.

Importantly, this way of modeling allowed us to directly compare several vaccination schedule specific parameters. We found that the Shingrix vaccination schedules led to a more pronounced proliferation of T-cells, however without a difference in T-cell decay rate between vaccination schedules. This shows the benefit of using of mathematical mixed models instead of performing a statistical analysis of the datasets: in the latter case it is possible to prove significant differences between vaccines, however, it is not determinable whether that increase is the result of either a higher proliferation of cells, a lower decay (mainly in the case of a restricted number of data points), or a longer time period (0, *h*) in which cells are activated.

We note that the adjuvant used for the Shingrix vaccine has been reported to be a very potent adjuvant [10] and our modeling approach thus confirms the increased proliferation of T-cells for the Shingrix vaccine.

We also assessed whether a correlation existed between the B-cell and T-cell counts, but we did not find a significant association between the two immune response types. This confirms previous findings concerning the glycoprotein-E adjuvant, part of the AS01_*B*_ Adjuvant System family, in which it has been shown that this family has been reported to show the lowest correlations between B-cells and T-cells of all families [11].

During our modeling analyses we encountered several limitations. First, we noted that given the limited sample size only models with moderate complexity could be analysed. Second, the sparseness of time points for the B-cell responses posed a significant limitation on the complexity of the B-cell models. Future work should focus on estimating an optimal sampling schedule for subsequent modeling. We conclude that nonlinear mixed modeling by means of ODE shows that Shingrix vaccination causes a significantly higher proliferation of T-cells compared to Varilrix vaccination in VZV-immune adults.

## Supporting information

**S1 Dataset Varilrix-specific B-cells**.

**S2 Dataset gE-specific B-cells**.

**S3 Dataset Varilrix-specific T-cells**.

**S4 Dataset gE-specific T-cells**.

**S1 Fig. Amount of VZV IgG-secreting cells** Measured by B-cell ELISPOT (gE stimulus) per 10^6^ IgG-secreting cells, up to 12 months. Data are shown per study group. The last panel shows a smooth function of the expected change in number of B-cells over time (in months), based on the observed data points per individual and considering the second vaccination at month 2.

**S2 Fig. Amount of gE-specific CD4+ T-cells, producing at least 2 immune markers**. easured by ICS per 10^6^ CD4+ T-cells, shown per group and up to 12 months. The last panel shows a smooth function of the expected change in number of T-cells over time (in months), based on the observed data points per individual.

**S3 Fig. Scatterplot of the increase in gE-specific T-cells (T01), plotted against the increase in gE-specific B-cells (B01)**.

**S1 Appendix. Detailed ODE models and solutions**. Antibody secreting cell models and T-cell models.

**S2 Appendix. Model selection of gE-specific B-cell data and T-cell data**.

**S3 Appendix. Algorithm parameter values used in Monolix**.

**S4 Appendix. Influence of the initial number of B-cells to the remaining B-cell data**.

## Acknowledgements

This research was funded by the University of Antwerp [BOF Concerted Research Action, Methusalem funding], NH acknowledges support from the chair in evidence-based vaccinology sponsored by a gift from Pfizer (2009-2018) and GSK (2016) and from the European Research Council (ERC) under the European Union’s Horizon 2020 research and innovation programme (grant agreement 682540 - TransMID). This research was supported by the Antwerp Study Centre for Infectious Diseases (ASCID). The funders had no role in study design, data collection and analysis, decision to publish, or preparation of the manuscript.

